# Evaluation of critical data processing steps for reliable prediction of gene co-expression from large collections of RNA-seq data

**DOI:** 10.1101/2021.03.11.435043

**Authors:** Alexis Vandenbon

## Abstract

**Motivation:** Gene co-expression analysis is an attractive tool for leveraging enormous amounts of public RNA-seq datasets for the prediction of gene functions and regulatory mechanisms. However, the optimal data processing steps for the accurate prediction of gene co-expression from such large datasets remain unclear. Especially the importance of batch effect correction is understudied.

**Results:** We processed RNA-seq data of 68 human and 76 mouse cell types and tissues using 50 different workflows into 7,200 genome-wide gene co-expression networks. We then conducted a systematic analysis of the factors that result in high-quality co-expression predictions, focusing on normalization, batch effect correction, and measure of correlation. We confirmed the key importance of high sample counts for high-quality predictions. However, choosing a suitable normalization approach and applying batch effect correction can further improve the quality of co-expression estimates, equivalent to a >80% and >40% increase in samples. In larger datasets, batch effect removal was equivalent to a more than doubling of the sample size. Finally, Pearson correlation appears more suitable than Spearman correlation, except for smaller datasets.

**Conclusion:** A key point for accurate prediction of gene co-expression is the collection of many samples. However, paying attention to data normalization, batch effects, and the measure of correlation can significantly improve the quality of co-expression estimates.

## Introduction

Understanding the functions and regulatory mechanisms of genes is one of the central challenges in biology. Gene co-expression is an important concept in bioinformatics because it serves as a foundation for predicting gene functions and regulatory mechanisms, and for more complex network inference methods (Eisen *et al*., 1998; Wolfe *et al*., 2005; Zhang and Horvath, 2005; Usadel *et al*., 2009; Serin *et al*., 2016; van Dam *et al*., 2018). Several gene co-expression databases have been developed (Zimmermann *et al*., 2004; van Dam *et al*., 2015; Vandenbon *et al*., 2016; Obayashi *et al*., 2019).

High numbers of samples are needed to accurately infer correlation of expression (Ballouz *et al*., 2015). Public databases are attractive sources of expression data, but in practice high numbers of samples can only be obtained by aggregating data from different studies conducted by different laboratories. As a result, the input data for gene co-expression analysis often contains considerable technical variability, called batch effects (Leek *et al*., 2010; Vandenbon *et al*., 2016).

In a recent study, we showed that correcting batch effects improved the quality of gene co-expression estimates significantly (Vandenbon *et al*., 2016). However, our previous study was limited to microarray data, and considered only one data normalization method and one batch correction method, i.e. ComBat (Johnson *et al*., 2007). Moreover, other studies have shown that treating batch effects can also result in unwanted artifacts such as exaggerated differences between covariates in gene expression and DNA methylation data (Harper *et al*., 2013; Nygaard and Rødland, 2016; Price and Robinson, 2018; Zindler *et al*., 2020).

Here, we present a systematic analysis of the effects of RNA-seq data normalization, batch effect correction, and correlation measure on the quality of gene co-expression estimates. We applied 50 data processing workflows on data for 68 human and 76 mouse cell types and tissues, resulting in 7,200 sets of genome-wide gene-gene co-expression predictions. Through analysis of the quality of these cell type- and tissue-specific co-expression predictions, we confirmed the importance of large numbers of samples (Ballouz *et al*., 2015). We also found that some normalization methods (especially UQ normalization) resulted on average in better co-expression predictions than others. In addition, treating batch effects resulted in a significant improvement of the co-expression estimates, especially in larger datasets consisting of samples produced by many different studies. It is imperative that future studies pay attention to batch effects in order to make optimal use of large amounts of public data. Finally, the difference between Pearson’s correlation and Spearman’s correlation was small, with Spearman working better in small datasets and Pearson better in medium-sized datasets. To the best of our knowledge, this is the first comprehensive study evaluating the importance of batch effect correction for the prediction of gene co-expression from large collections of RNA-seq data.

## Results

### Overview of this study

The goal of this study is to gain insights into which data processing steps are preferable for obtaining high-quality gene co-expression estimates from large collections of RNA-seq data. To address this issue, we collected a dataset of 8,796 human and 12,114 mouse bulk RNA-seq samples, from 401 and 630 studies, covering 68 human and 76 mouse cell types and tissues (see Methods; Supplementary Tables S1 and S2) (Petryszak *et al*., 2017). On these two datasets, we applied combinations of data normalization approaches and batch effect correction approaches (see Figure 1 for a summary of the workflow). As proxies for batches we used the studies that produced each sample (1 study is 1 batch). We also applied the method ComBat-seq on the raw read count data without any prior normalization (Zhang *et al*., 2020). The resulting 25 (6 normalizations x 4 batch effect correction approaches, and ComBat-seq without normalization) human and 25 mouse datasets were used to estimate correlation of expression using Pearson’s correlation and Spearman’s correlation in the data of each cell type or tissue. This resulted in a total of 7,200 (3,400 human and 3,800 mouse) genome-wide sets of cell type or tissue-specific gene co-expression predictions. We will refer to each genome-wide set of cell type or tissue-specific gene co-expression predictions as a “co-expression network”. However, the goal of this study is not to analyze network topology. Our focus is to identify the key features that result in accurate co-expression predictions.

**Figure 1:**
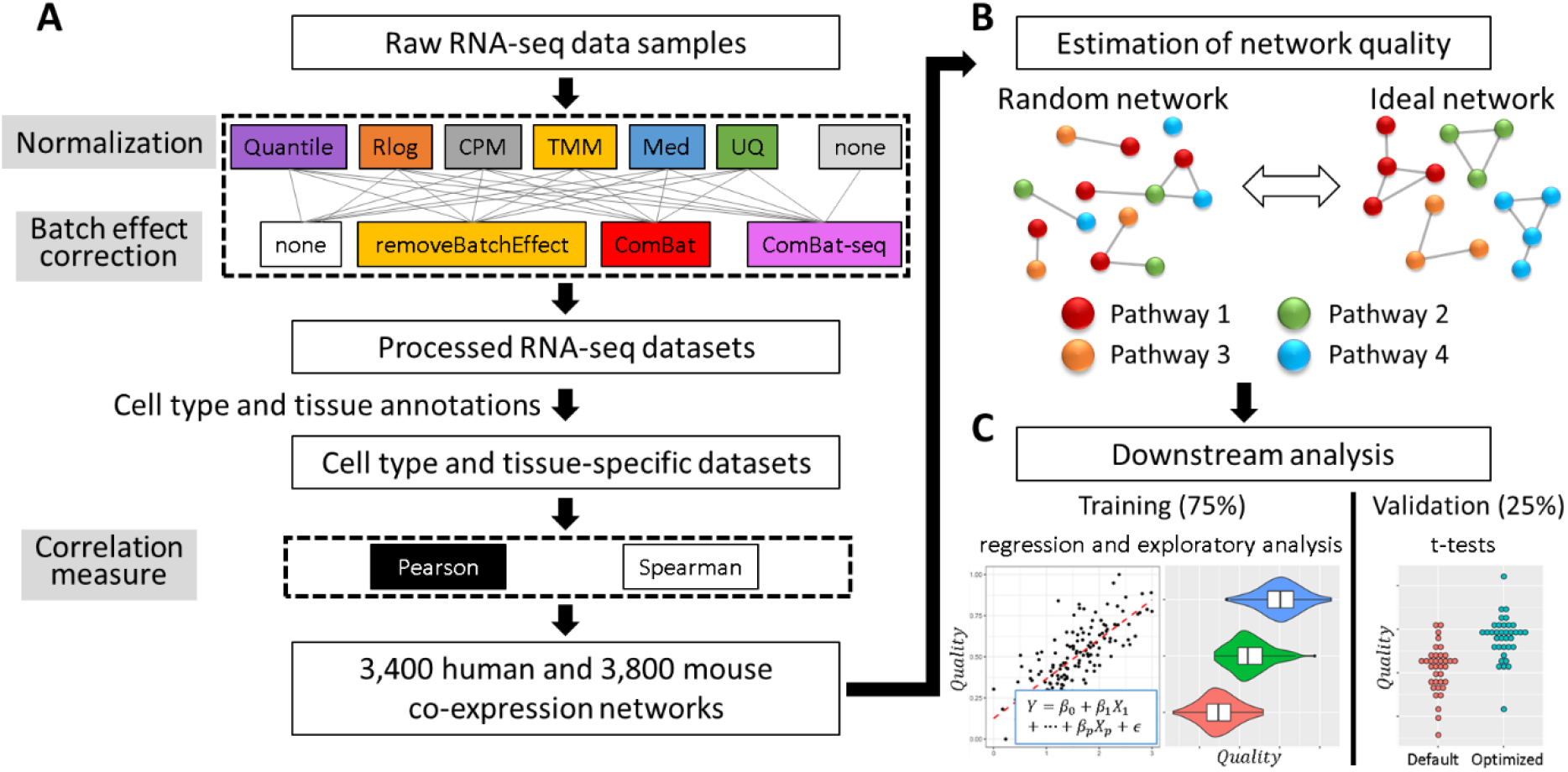
Summary of this study. **(A)** Raw RNA-seq data was processed with 50 different combinations of normalization, batch effect correction, and correlation measures into 7,200 genome-wide sets cell type and tissue-specific co-expression predictions, which we refer to as “co-expression networks”. **(B)** Quality of co-expression networks was estimated based on the enrichment of functional annotations of correlated genes and regulatory motifs in their promoters. In random co-expression networks no common annotations and motifs are expected to be found among correlated genes. In contrast, in ideal networks such enrichments should be encountered frequently. Here nodes represent genes and edges co-expression. **(C)** Quality measures were processed into 7,200 quality scores, which were used for downstream analysis. We use 75% of cell types and tissues for regression and exploratory analysis, and the remaining 25% for validation.

### Defining the quality of co-expression predictions

Next, we evaluated the quality of the co-expression predictions produced by each workflow. Many studies have used enrichment of shared functional annotations among correlated genes or regulatory DNA motifs in their promoter sequences as quality measures for co-expression predictions (Lee *et al*., 2004; Verleyen *et al*., 2015; Ballouz *et al*., 2017; Obayashi *et al*., 2019). In high-quality co-expression networks, we expect correlated genes to belong to shared pathways or to be controlled by a common regulatory mechanism (Figure 1B). In contrast, in a randomly generated network, correlated genes are expected to lack common functions or regulatory mechanisms. In this study, in each co-expression network, for every gene, we extracted the set of 100 genes with the highest correlation. We then defined eight quality measures that are based on how frequently we observed significant enrichment of functional annotations (GO terms) and regulatory motifs (in promoter sequences) among these sets of 100 genes (see Supplementary Methods for more details). In high-quality networks this frequency should be high, and in low-quality networks it should be low.

We collected the eight quality measures of the 7,200 co-expression networks and Principal Component Analysis (PCA) revealed that they are highly consistent and correlated: the first PC explained 81.4% of variability in the quality measures (Supplementary Figure S1A), and had a high correlation with all eight measures (range 0.77 to 0.96; Supplementary Figure S1B-C). To facilitate the downstream analysis, we therefore decided to use this first PC as a general quality score (*Quality*, see Methods) after rescaling it to the range 0 (worst networks) to 1 (best networks) (Supplementary Figure S1D). As an illustration, Supplementary Figure S2 shows the quality measures for the networks with the lowest (Med + ComBat-seq + Spearman applied on human salivary gland data), 50^th^ percentile (Rlog + ComBat + Pearson applied on human neuron data), and highest (UQ + removeBatchEffect + Spearman applied on mouse liver data) *Quality*. From the lowest-quality network to the highest-quality network, the measures of quality are progressively increasing.

At this point we randomly split the 68 human and 76 mouse cell types and tissues into 2 groups (Supplementary Tables S1 and S2). One group (51 human and 57 mouse cell types and tissues) will be used for the analysis of features that contribute to high-quality gene co-expression predictions. We refer to this as our training set, and will focus on it in the following sections. The other group (17 human and 19 mouse cell types and tissues) will be used as an independent validation set later (section “The best workflows result in significantly better co-expression estimates”).

Figure 2 shows the 50 workflows that we examined, sorted by the average *Quality* of the 108 (51 human and 57 mouse cell types and tissues in the training set) networks that they each resulted in. We observed that the top four workflows used UQ normalization, while Quantile normalization resulted in low average quality. Similarly, the top 15 workflows all include a batch correction step, while many of the worst-performing workflows did not treat batch effects. The top-ranking workflow (UQ + ComBat + Pearson) resulted in an above-average network for 102 (94%) of the 108 training datasets, and for 135 (94%) of all 144 datasets (Supplementary Figure S3).

**Figure 2:**
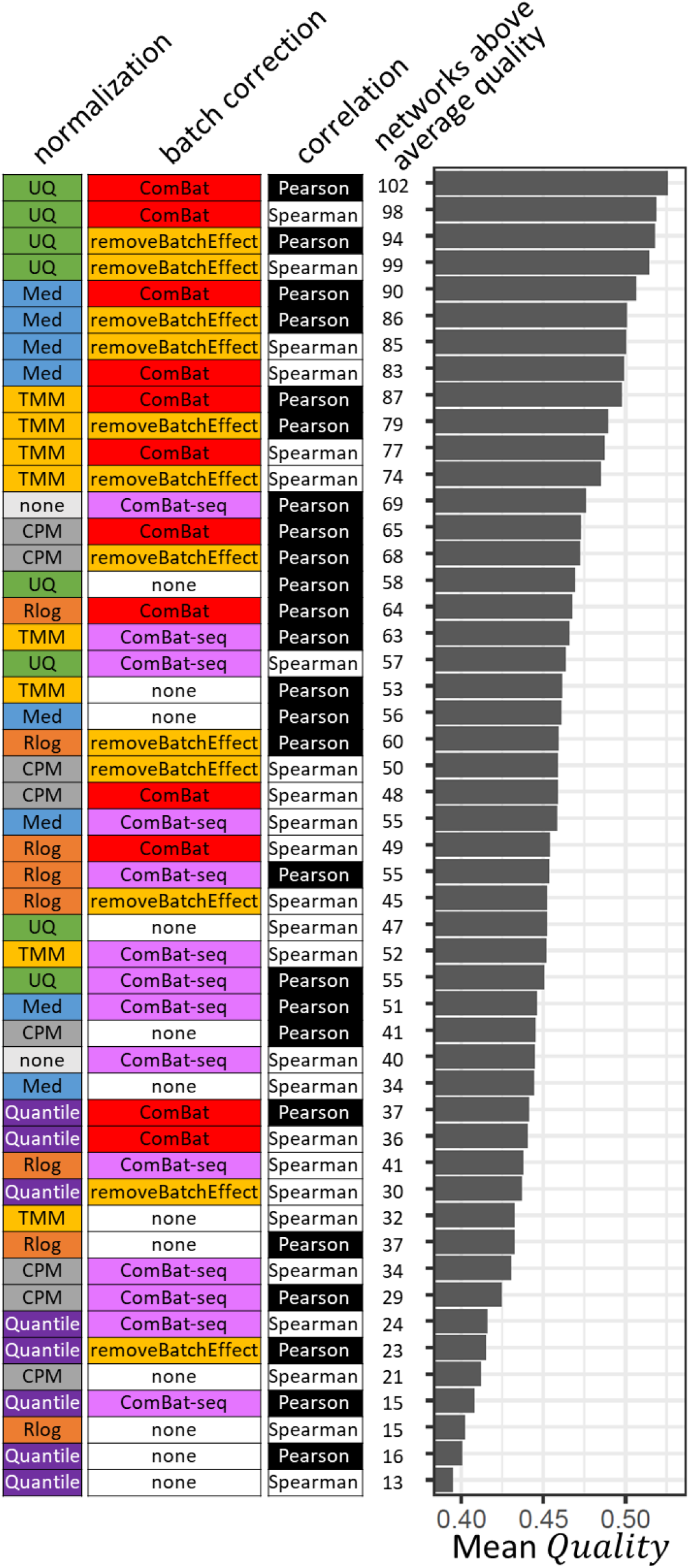
Evaluating the quality of co-expression networks. All 50 workflows are shown in order of decreasing average quality of the networks they produced. From left to right are shown: normalization method, batch effect correction method, and measure of correlation used in each workflow. Next, the number of training datasets (108 in total) in which the workflow resulted in an above-average quality network is shown, and the mean *Quality* of the 108 networks generated using each workflow.

### Modeling the quality of co-expression networks

To gain more quantitative insights into what factors contribute to high-quality co-expression estimates, we performed linear regression on the *Quality* scores using as predictors: 1) the number of RNA-seq samples which the network was based on (log_10_ values), 2) the number of batches in the data (log_10_values), 3) the species (human or mouse), 4) the data normalization approach, 5) batch correction approach, and 6) the correlation measure. In the next sections, we will focus on workflows that did not include ComBat-seq. ComBat-seq differs from ComBat and RemoveBatchEffect in that it takes integers as input and therefore cannot be used on data that has already been normalized. Networks generated using ComBat-seq will be treated separately in section “ComBat-seq results in lower-quality networks”.

The resulting linear model is summarized in Table 1. Despite its simplicity, this model explains 55% of the variability in *Quality* (R^2^ = 0.55) in the training datasets. Below we discuss the roles of sample counts, data normalization, batch effect correction, and correlation measures in more detail.

**Table 1:**
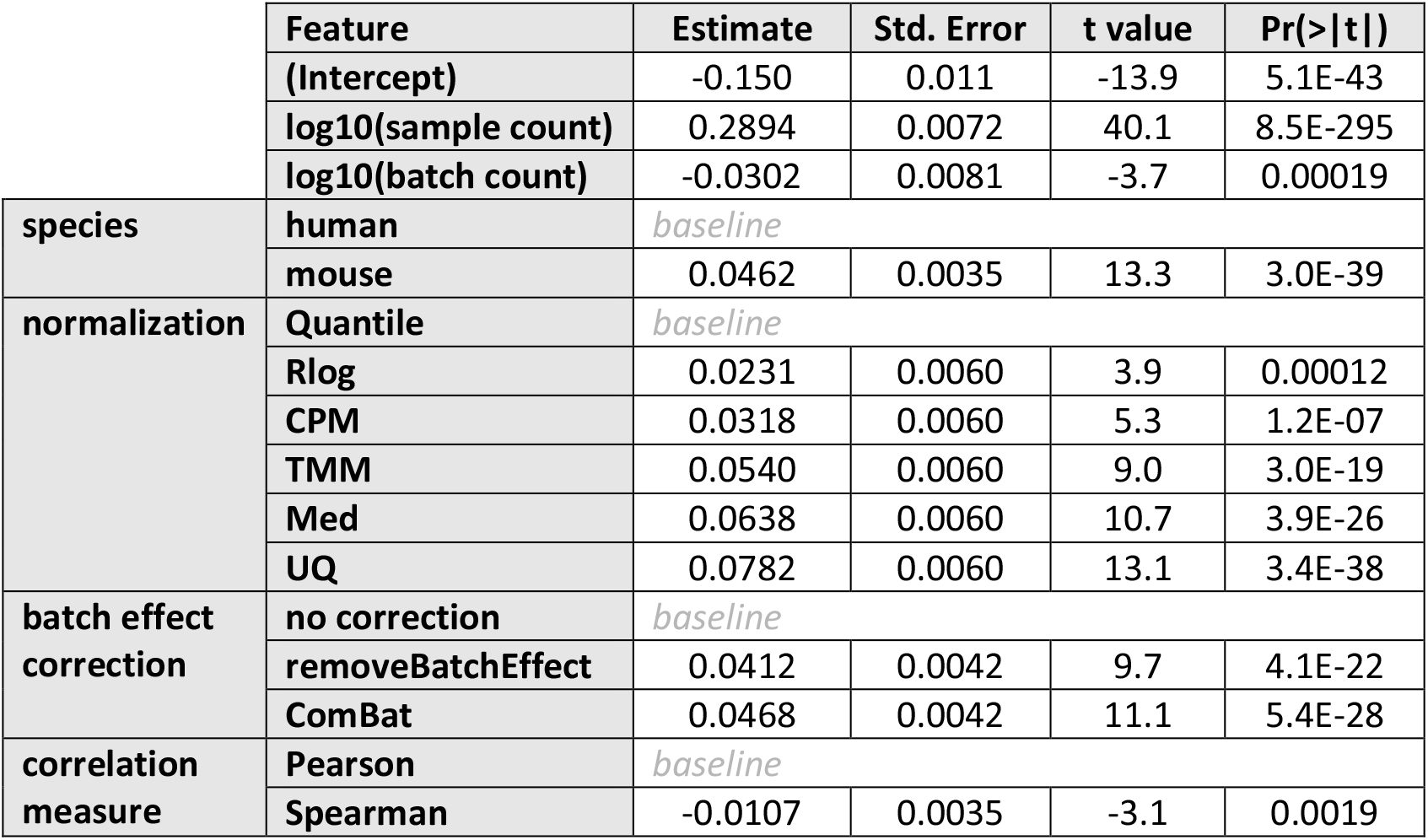
Linear regression analysis of co-expression network quality scores. Features, their estimated coefficient, standard error, t value and p-value are shown. Qualitative predictors are grouped by species, normalization, batch effect correction and correlation measure.

|  | Feature | Estimate | Std. Error | t value | Pr(>\|t\|) |
| --- | --- | --- | --- | --- | --- |
|  | (Intercept) | -0.150 | 0.011 | -13.9 | 5.1E-43 |
|  | log10(sample count) | 0.2894 | 0.0072 | 40.1 | 8.5E-295 |
|  | log10(batch count) | -0.0302 | 0.0081 | -3.7 | 0.00019 |
| species | human | baseline |  |  |  |
|  | mouse | 0.0462 | 0.0035 | 13.3 | 3.0E-39 |
| normalization | Quantile | baseline |  |  |  |
|  | Rlog | 0.0231 | 0.0060 | 3.9 | 0.00012 |
|  | CPM | 0.0318 | 0.0060 | 5.3 | 1.2E-07 |
|  | TMM | 0.0540 | 0.0060 | 9.0 | 3.0E-19 |
|  | Med | 0.0638 | 0.0060 | 10.7 | 3.9E-26 |
|  | UQ | 0.0782 | 0.0060 | 13.1 | 3.4E-38 |
| batch effect correction | no correction | baseline |  |  |  |
|  | removeBatchEffect | 0.0412 | 0.0042 | 9.7 | 4.1E-22 |
|  | ComBat | 0.0468 | 0.0042 | 11.1 | 5.4E-28 |
| correlation measure | Pearson | baseline |  |  |  |
|  | Spearman | -0.0107 | 0.0035 | -3.1 | 0.0019 |

### The importance of sample counts

The most significant predictor for the quality of co-expression estimates was the number of samples they were based on (Table 1). *Quality* follows a roughly linear trend with the logarithm of the sample count (Figure 3A). This is consistent with a previous study (Ballouz *et al*., 2015). At the same time, an increase in the number of batches results in a small decrease in *Quality* (Table 1). A number of samples obtained from a small number of batches is expected to result in better co-expression predictions than an equal number of samples generated by many smaller batches.

**Figure 3:**
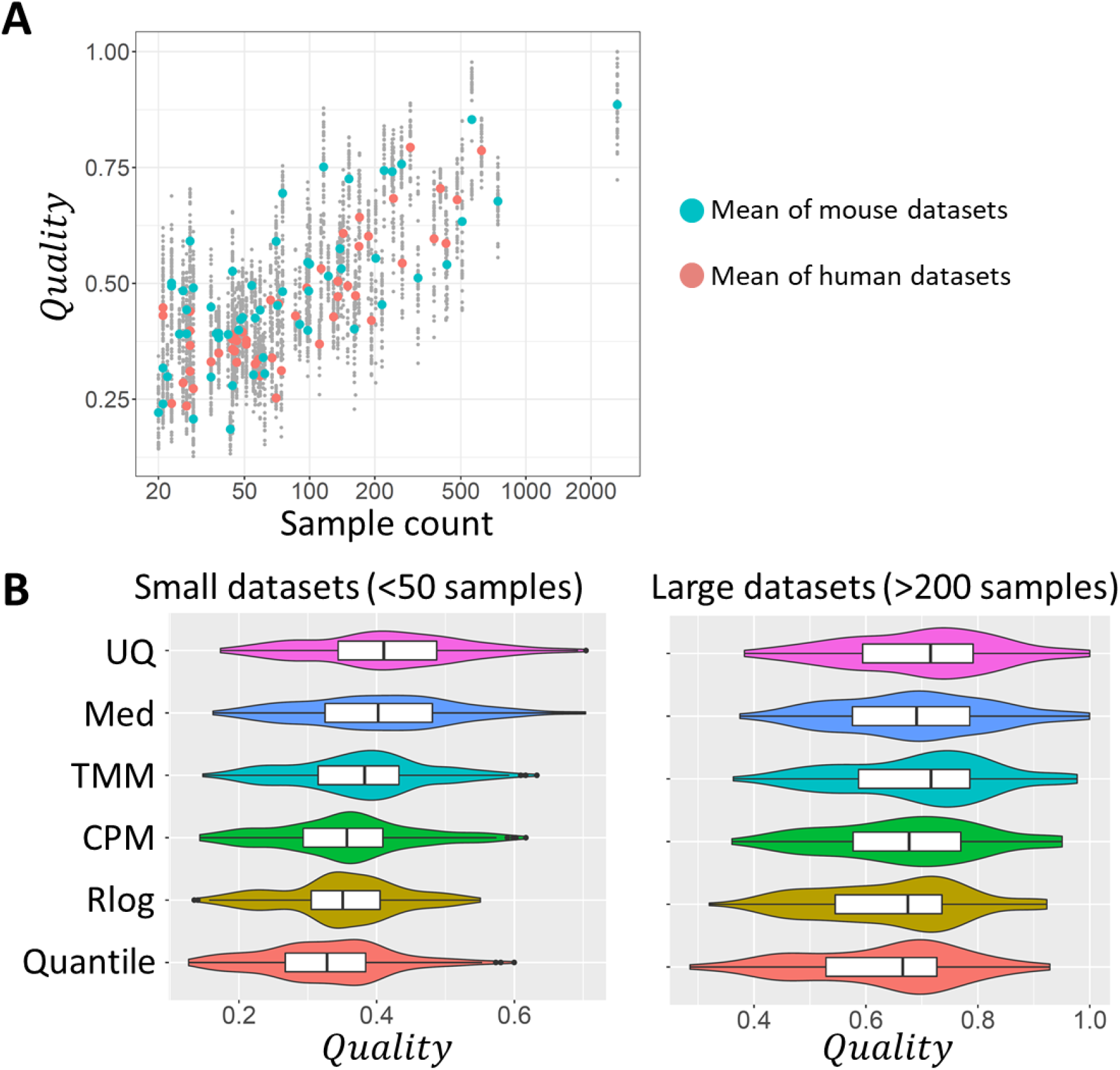
Importance of sample numbers and normalization approaches. **(A)** Sample count vs quality of co-expression networks. Small points represent the individual networks. Larger points are averages for each dataset. Blue: mouse, red: human datasets. **(B)** Violin plots of the quality of networks made using each of the six normalization methods. Left: for 44 small datasets (< 50 samples). Right: for 19 large datasets (> 200 samples).

*Quality* being roughly linearly related to the logarithm of the sample count implies that an ever-increasing number of additional samples is needed to achieve the same improvement in quality. Collecting hundreds or thousands of additional samples is practically impossible under most circumstances. Therefore, it makes sense to look for data processing steps that can maximize the quality of co-expression predictions even in the absence of an increase in samples.

### The importance of data normalization approach

Regression analysis revealed clear differences in the average quality of networks generated using the six different normalization methods (Table 1), confirming the tendencies observed in Figure 2. Med and UQ resulted in an average increase of 0.064 and 0.078 in *Quality* compared to the baseline (here Quantile normalization, the worse performing method), respectively. These improvements are equivalent to a 66% and 86% increase in sample count. Additional exploratory analysis revealed interactions exist between sample counts and normalization methods: the performance of normalization methods depends on the size of datasets. Figure 3B shows *Quality* of networks based on small datasets (< 50 samples) and large datasets (> 200 samples). UQ performed well on both large and small datasets. In contrast, TMM performed well on large datasets, but less so on small datasets (Figure 3B).

### Correcting batch effects in general improves co-expression quality

Correction of batch effects by removeBatchEffect or ComBat resulted in better networks, increasing the *Quality* on average by respectively 0.041 and 0.047 compared to no correction (Table 1). These improvements are equivalent to a 39% and 45% increase in sample count, respectively. However, here too, the improvement depends on sample counts, and on the number of batches in the dataset. The improvement in quality appears to increase roughly with the sample count, for both removeBatchEffect (Figure 4A) and ComBat (Figure 4B). Especially ComBat consistently resulted in higher-quality networks in larger datasets (Figure 4A-B). For datasets with >200 samples, ComBat’s average improvement in *Quality* exceeded 0.10, equivalent to a >120% increase in sample count. removeBatchEffect failed to correct batch effects in a few datasets, resulting in somewhat worse overall quality (Figure 4A, indicated datasets). Batch effect correction offered no clear advantage when a dataset contained few batches (e.g. less than 5, Figure 4D-E). However, for datasets containing 5 or more batches, using ComBat or removeBatchEffect resulted in general in better networks.

**Figure 4:**
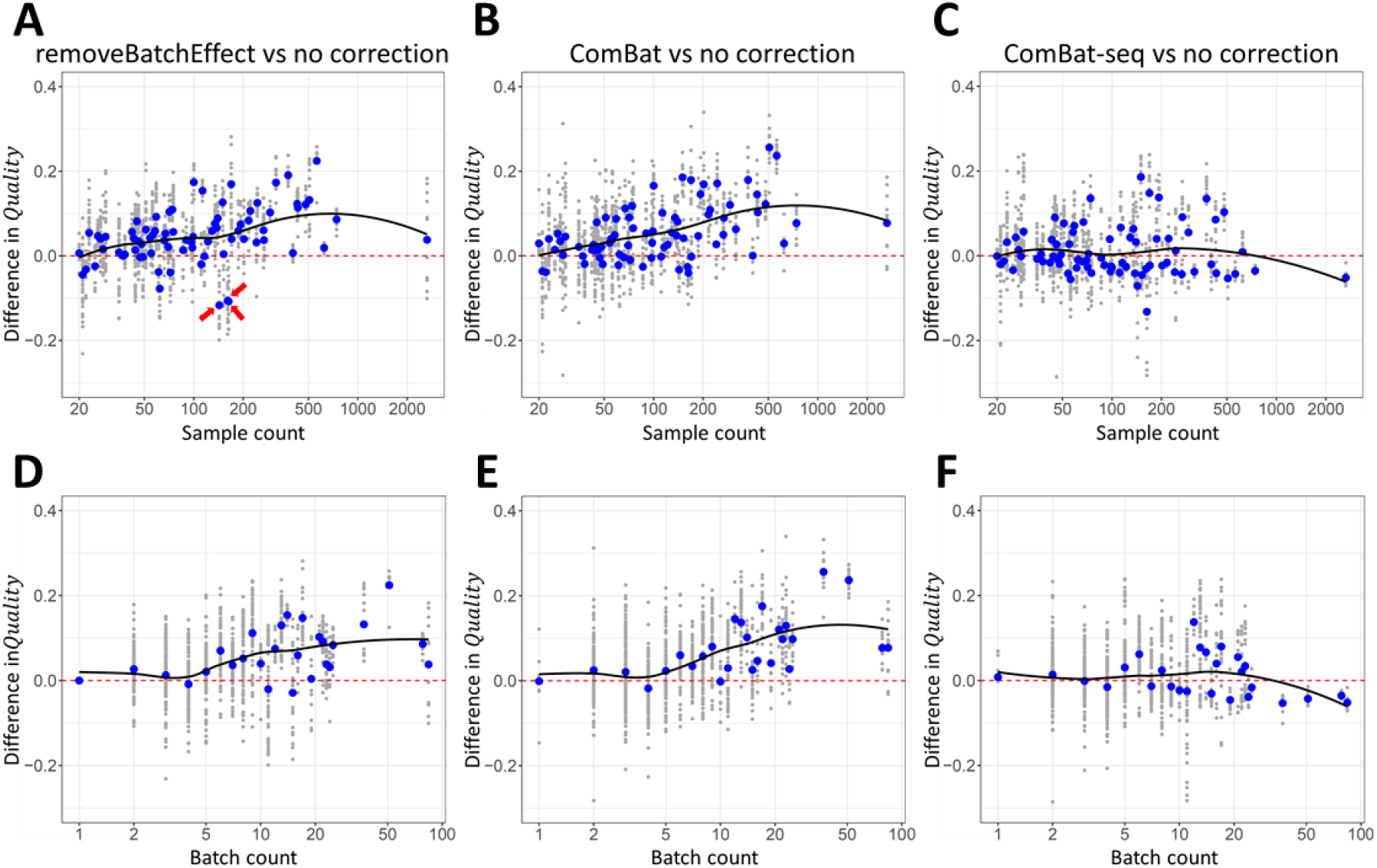
The number of samples and batches affect the advantage of batch effect correction. **(A-C)** Difference in quality of co-expression networks based on data with and without treating batch effects in function of sample count, for data treated with removeBatchEffect **(A)**, ComBat **(B)**, and ComBat-seq **(C)**. Positive values indicate an increase in quality in the batch-treated networks. **(D-F)** The same differences in quality in function of the number of batches in each dataset, for removeBatchEffect **(D)**, ComBat **(E)**, and ComBat-seq **(F)**. The average difference in quality is shown in blue for each sample count and for each batch count. A smoothed pattern (loess) is included in each plot.

### Spearman correlation is preferred for small datasets

Using Spearman’s correlation instead of Pearson correlation resulted on average in a 0.011 decrease in *Quality* (equivalent to a 8.2% decrease in sample count) (Table 1). However, for small datasets (sample count < 30) Spearman’s correlation had on average an advantage (Figure 5). For medium-sized datasets (roughly 30 to 100 samples) Pearson’s correlation lead in general to better co-expression networks, but the difference became smaller with higher sample counts.

**Figure 5:**
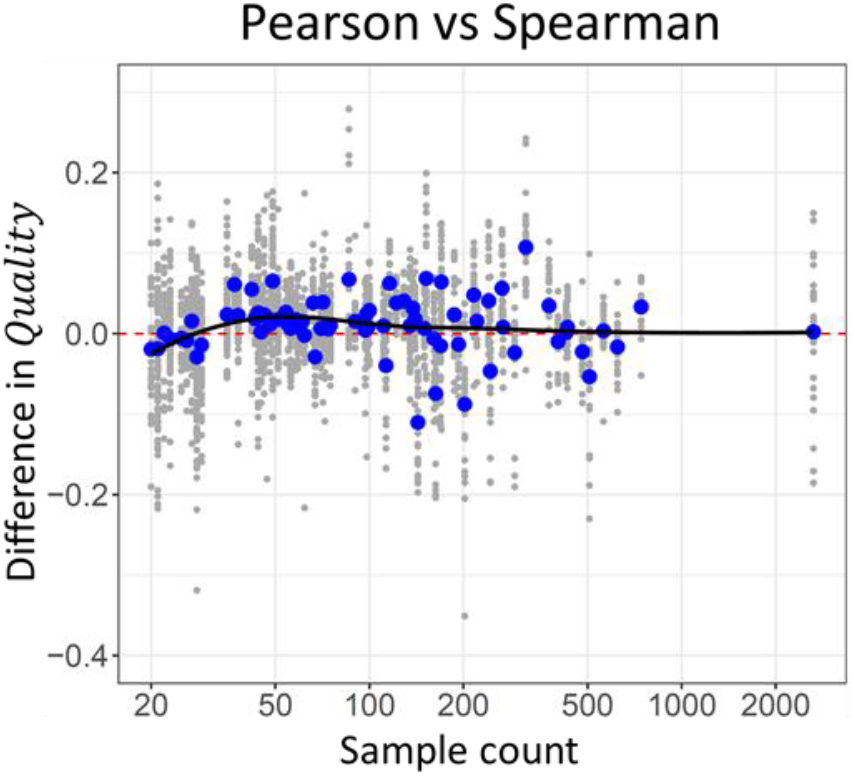
The preferred correlation measure depends on the number of samples. Difference in the quality of co-expression networks based on Pearson’s correlation and networks based on Spearman’s correlation is shown in function of the number of samples in the datasets. Positive values indicate an advantage for Pearson’s correlation. The average difference in quality is shown in blue for each dataset. A smoothed pattern (loess) is included in the plot.

These results make intuitive sense. Spearman’s correlation, which is based on ranks and not on raw values, is less sensitive to extreme values. In small datasets, extreme values have a strong influence on correlation, adversely affecting Pearson’s correlation. However, in medium-sized datasets, the influence of extreme values decreases and Pearson’s correlation appears to be better able to capture biological signals.

### ComBat-seq results in lower-quality co-expression estimates

On average, ComBat-seq did not result in high-quality networks compared to ComBat and removeBatchEffect (Figure 2). Adding a normalization step following the correction by ComBat-seq did not improve qualities but rather reduced them (Figure 2). Zhang and colleagues themselves noted that on some datasets ComBat-seq did not outperform ComBat (Zhang *et al*., 2020). In our data, ComBat-seq resulted in lower-quality networks even in the datasets with more samples (Figure 4C) or more batches (Figure 4F). Although the reason for this failure is not clear, we did note that ComBat-seq returned unrealistically high read counts in a substantial subset of samples. For example, in 821 human samples (out of a total of 8,796) the total read count after correction exceeded 10 billion reads. A subset of genes had extremely high reads counts in some samples. For example, *Prh1* had corrected read counts > 1e100 in a subset of human salivary gland samples. These observations suggest that ComBat-seq adjusted a subset of the data to negative binomial distributions that are extremely skewed, negatively affecting the quality of co-expression networks.

### The best workflows result in significantly better co-expression estimates

Finally, we returned our attention to the eight raw quality measures, and the validation datasets. We compared the networks produced by a default workflow (Rlog + no batch correction + Pearson) with those produced by optimized workflows (UQ + ComBat + Spearman for datasets with < 30 samples, and UQ + ComBat + Pearson otherwise) (Figure 6). For all eight measures, using the optimized approaches resulted in a significant improvement compared to the default workflow (one-sided paired t-tests; all eight p-values < 0.01; improvements seen in 25 to 33 of the 36 validation datasets). The optimized workflows lead to co-expressed genes sharing common functional annotations more frequently (Figure 6A-C left side), as well as shared annotations fitting with the known annotations of genes more frequently (Figure 6A-C right side). The same was true for regulatory motifs in promoter sequences (Figure 6D).

**Figure 6:**
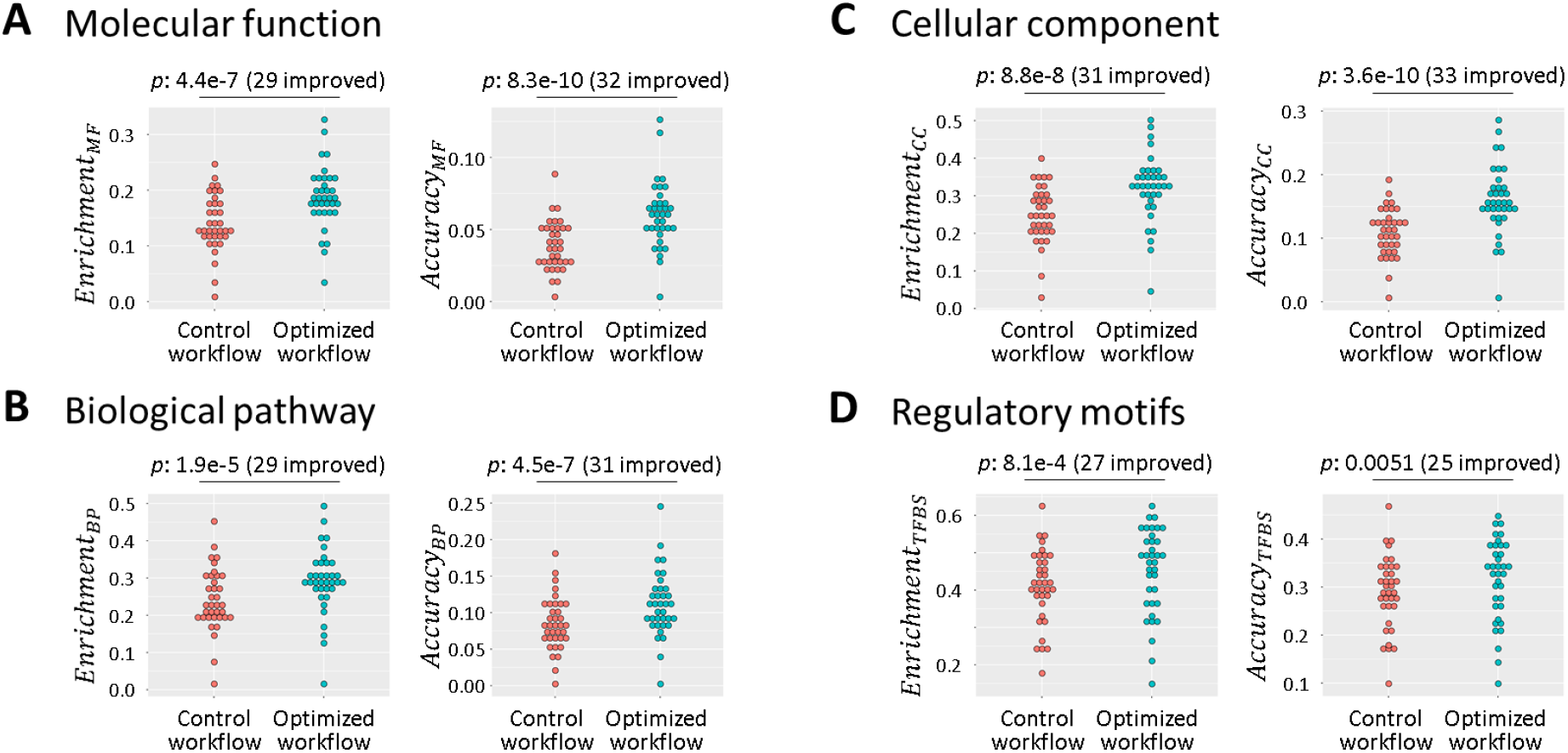
Optimized workflows lead to a significant improvement in all eight quality measures. **(A-D)** (left) Dotplots showing the fraction of genes with enrichment of GO terms and regulatory motifs in networks produced by a default workflow (red) and the optimized workflow (blue). (right) Dotplots showing the fraction of genes with an annotation that fit with enrichment GO terms, and with promoters that contain an instance of an enriched regulatory motif. Each dot represents one of the 36 (17 human and 19 mouse cell types and tissues) validation datasets. P values are based on one-sided paired t-tests. The number of validation datasets in which an improvement was observed is indicated between brackets.

## Discussion

We presented a systematic analysis of 50 workflows for processing large collections of RNA-seq data into gene co-expression predictions. We applied the workflows on data for 68 human and 76 mouse tissues and cell types, and estimated the quality of the resulting 7,200 sets of genome-wide gene co-expression datasets (“co-expression networks”). We used linear regression analysis to gain understanding of the important factors for obtaining high-quality co-expression networks. Our aim was to re-analyze existing large RNA-seq expression datasets, that have already been trimmed, aligned to a reference genome, and counted using a standardized pipeline. We focused on the steps of RNA-seq data normalization, batch effect correction, and measure of correlation of expression. Other studies have compared between read trimming and alignment approaches in related contexts (Corchete *et al*., 2020).

We found that co-expression network quality is to a large degree determined by the number of samples on which it is based, as has been reported before (Ballouz *et al*., 2015). It is therefore important to gather as many samples as possible. However, in practice the number of available samples is always limited, so it makes sense to optimize the data processing workflow to obtain high-quality networks even in the absence of large sample counts.

Treating batch effects in general lead to better networks. On average, ComBat performed better than limma’s removeBatchEffect function. ComBat-seq however performed considerably worse than ComBat and removeBatchEffect. In our analysis, correcting batch effects using ComBat resulted in improvements to network quality equivalent to a 45% increase in sample count on average. In larger datasets, the advantage was even more pronounced, equivalent to roughly a doubling in sample count. Unfortunately, gene co-expression studies still often ignore batch effects. Clearly, more attention needs to be paid to the issue of batch effects in order to extract the maximum potential out of ever-increasing public gene expression datasets.

We found that some data normalization approaches lead to better co-expression estimates than others. Especially UQ performed well. UQ has also been found to perform relatively well compared with total count normalization (equivalent to CPM in our study) and quantile normalization in the context of predicting differentially expressed genes (Bullard *et al*., 2010). Other comparisons of RNA-seq normalization methods (outside of the context of co-expression) have come to different and conflicting conclusions (Dillies *et al*., 2013; Li *et al*., 2015; Conesa *et al*., 2016).

The measure of correlation appeared to be less crucial, but Pearson’s correlation seems to have a slight advantage, except when there are only small numbers of samples (<30). In the latter case, Spearman’s correlation seems better.

Although no workflow dominated all others, UQ + ComBat + Pearson (or Spearman for small datasets) resulted in the best quality overall, and in above-average co-expression networks in >90% of the tissues and cell types we examined.

## Methods

### Gene expression data collection and normalization

We used the RNASeq-er REST API of the European Bioinformatics Institute (Petryszak *et al*., 2017) to obtain read count data for 8,796 human and 12,114 mouse RNA-seq samples, produced by 401 and 630 studies, covering 68 human and 76 mouse cell types and tissues, respectively (see Supplementary Methods and Supplementary Tables S1 and S2). On these two large datasets of human and mouse samples, we applied the following six normalization approaches:

#### Trimmed Mean of M-values (TMM)

For all genes, log ratios are calculated versus a reference sample (Robinson and Oshlack, 2010). The most highly expressed genes, and genes with high log ratios are filtered out. The mean of the remaining log ratios is used as a scaling factor. This normalization method is the default normalization method of the edgeR function calcNormFactors (McCarthy *et al*., 2012; Robinson and Oshlack, 2010). In this study, we first removed genes that have less than 1 read per million reads in all samples prior to normalizing the remaining genes.

#### Counts per million (CPM)

the number of reads per gene is divided by the total number of mapped reads of the sample and multiplied by 1 million (Dillies *et al*., 2013; Abbas-Aghababazadeh *et al*., 2018).

There are several variations on CPM, describe below, including Upper Quartile and Median.

#### Upper Quartile (UQ)

counts are divided not by total count but by the upper quartile of non-zero values of the sample (Bullard *et al*., 2010).

#### Median (Med)

counts are divided not by total count, but by the median of non-zero values of the sample (Dillies *et al*., 2013).

#### Regularized Logarithm (RLog or RLE)

A regularized-logarithm transformation is applied which is similar to a default logarithmic transformation, but in which lower read counts are shrunken towards the genes’ averages across all samples. We applied this normalization using the R package DESeq2 (Love *et al*., 2014).

#### Quantile

All samples are normalized to have the same quantiles. We applied this normalization using the function normalizeQuantiles of the limma R package (Ritchie *et al*., 2015).

Note that methods that correct for differences in gene length (RPKM and FPKM) are not relevant here, since they don’t affect correlation values. In this study, these methods would be equivalent to CPM normalization.

We thus obtained 12 normalized datasets (6 each for the human and the mouse data). Each dataset was transformed to log values after addition of a small pseudo count (defined as the 1% percentile of non-zero values in the normalized dataset).

### Batch effect correction using Combat and limma’s removeBatchEffect function

On the 12 log-transformed datasets we applied two batch effect correction methods: ComBat (function ComBat in the sva R package) and the removeBatchEffect function of the limma R package (Johnson *et al*., 2007; Ritchie *et al*., 2015). Both ComBat and removeBatchEffect allow users to specify batch covariates to remove from the data (“batch” parameter in both functions). Here, studies were used as substitutes for batches. Users can also give biological covariates to retain (“mod” parameter in ComBat and “design” parameter in removeBatchEffect). In this study, biological covariates are the cell type or tissue from which the samples were obtained. Both Combat and removeBatchEffect were used using default parameter settings.

To be able to treat batch effects in a dataset there can be no confounding between technical and biological covariates. In practice, studies often focus on a single cell type, which makes confounding of batches and cell types highly probable. Both the human and mouse datasets could be divided into several subsets with no shared cell types or tissue annotations. Therefore, batch effects were corrected for each of these subsets of samples separately, and finally the treated datasets were merged again into one dataset.

In addition to correcting the data using ComBat and removeBatchEffect we also considered 2 other options. One is to ignore batch effects and use the normalized data directly for estimating gene co-expression. Another is to use ComBat-seq.

### Batch effect correction and normalization using Combat-seq

ComBat-seq differs from ComBat and limma’s removeBatchEffect function in that both its input and output are integer counts, making it more suitable for RNA-seq read count data (Zhang *et al*., 2020). To correct batch effects using ComBat-seq (function Combat_seq in the sva R package), we therefore gave as input the raw count data (without any normalization step applied to it), as well as the same batch covariates and biological covariates as we used for ComBat and removeBatchEffect. Combat-seq was used with default parameter settings. The output read counts were transformed to log values after adding a pseudocount of 1. These log-transformed data were used for estimating correlation of expression (see next section) without an additional normalization step.

However, quality estimates of the co-expression networks generated using ComBat-seq were relatively low compared to ComBat and removeBatchEffect, which use normalized data as input (Figure 2 and Figure 4C,F). Therefore, to avoid an unfair comparison, we also applied the 6 normalization approaches (TMM, CPM, UQ, Med, Rlog and quantile) on the ComBat-seq output data. Rlog normalization failed because of the extremely high reads counts in a subset of the data (see also section “ComBat-seq results in lower-quality co-expression estimates”). We therefore trimmed all read counts >1e9 to 1e9 before conducting the Rlog normalization.

Together, this resulted in 7 datasets which had been processed using ComBat-seq (i.e. 6 with a normalization step, and 1 without).

### Estimating Correlation of Expression

For each of the 25 data processing combinations (6 normalization methods x 4 batch correction methods, and ComBat-seq without normalization), we calculated correlation of expression between each pair of genes in the log-transformed data for each cell type or tissue. We did this using Pearson’s correlation and Spearman’s rank correlation. Before calculating correlation coefficients in the expression data of a tissue or cell type, genes with general low levels of expression (less than 10 mapped reads in >90% of samples) or with no variation in expression (standard deviation = 0) were removed. The number of samples and genes used for calculating co-expression in each tissue and cell type are listed in Supplementary Tables S1 and S2.

### Evaluation of gene co-expression network quality

The processing steps described above resulted in 7,200 (3,400 human and 3,800 mouse) sets of genome-wide cell type or tissue-specific gene co-expression predictions, which we refer to as “co-expression networks”. The main goal of this study is to gain understanding into what are the critical features that distinguish good co-expression networks from bad ones. Because there is no gold standard co-expression network available, we first defined eight measures of quality based on the enrichment of biologically meaningful features (functional annotations of genes and regulatory DNA motifs in promoter sequences) among co-expressed genes. To facilitate comparison between co-expression networks, these eight measures were finally combined into a single quality score, *Quality*, with values ranging between 0 (worst) and 1 (best). We refer to Supplementary Methods for a more detailed description.

### Linear regression analysis

We conducted least squares regression using the lm function in R. As response variable we used *Quality*, and as predictors we used 1) the number of RNA-seq samples on which the co-expression network was based (log_10_ values), 2) the number of batches in the data (log_10_ values), 3) the species (human or mouse), 4) the data normalization approach, 5) batch correction approach, and 6) the correlation measure. For the categorical predictors (i.e. species, data normalization approach, batch correction approach and correlation measure) the lm function uses dummy variables with values 0 and 1. The baseline levels of these categorical variable were set in a way that facilitates interpretation of the results.

## Data availability

Both the human and the mouse RNA-seq datasets are available in figshare with DOIs doi.org/10.6084/m9.figshare.14178446.v1 and doi.org/10.6084/m9.figshare.14178425.v1. These datasets include 1) raw read counts of genes in all RNA-seq samples, 2) the same RNA-seq data after UQ normalization and batch effect correction using ComBat, which is in general the best workflow according to our study, 3) annotation data assigning a study ID and cell type or tissue to each sample, and 4) A list of each cell type or tissue included in the dataset along with its sample count.

## End notes

### Author contributions

A.V. conceived of the project and methodology, ran the analyses, and wrote the manuscript.

## Acknowledgements

We thank Dr. Diego Diez (Osaka University), Prof. Yoshio Koyanagi and the members of the Lab. of Systems Virology (Kyoto University), Prof. Kenta Nakai and the member of the Lab. of Functional Analysis in silico (Tokyo University), Prof. Wataru Fujibuchi (Kyoto University) for helpful discussions and advice. This work was supported by a KAKENHI Grant-in-Aid for Scientific Research (C) (20K06609) by the Japan Society for the Promotion of Science (JSPS).

