## Supplementary Material for "Evaluation of critical data processing steps for reliable prediction of gene co-expression from large collections of RNA-seq data"

Alexis Vandenbon[1,2]

[1] Institute for Frontier Life and Medical Sciences, Kyoto University, 53 Shougoin Kawara-cho, Sakyou-ku, Kyoto 606-8507, Japan

[2] Institute for Liberal Arts and Sciences, Kyoto University, Yoshidanihonmatsu-cho, Sakyo-ku, Kyoto 606-8501, Japan

**Contact:**

### Supplementary Methods

#### Gene Expression Data

We used the RNASeq-er REST API of the European Bioinformatics Institute (EBI; https://www.ebi.ac.uk/fg/rnaseq/api/, (Petryszak *et al.*, 2017)) to obtain a list of human and mouse RNA-seq datasets from the European Nucleotide Archive (ENA) with at least 70% of reads mapped to their reference genome, using URLs:

http://www.ebi.ac.uk/fg/rnaseq/api/tsv/70/getRunsByOrganism/homo_sapiens

http://www.ebi.ac.uk/fg/rnaseq/api/tsv/70/getRunsByOrganism/mus_musculus

Annotation data for each study was downloaded (http://www.ebi.ac.uk/fg/rnaseq/api/tsv/getSampleAttributesPerRunByStudy/<study ID>), and parsed for "cell type" and "organism part" annotation fields. Annotation terms were manually checked and turned into a list of consistent terms (removing differences and redundancies in spelling and capitalization). This resulted in consistent cell type or tissue annotations for the RNA-seq samples.

For all samples to which we could assign an annotation, raw read counts per gene were downloaded using the URL:

ftp://ftp.ebi.ac.uk/pub/databases/arrayexpress/data/atlas/rnaseq/studies/ena/<study ID>/<organism>/genes.raw.tsv

where <organism> is either "mus_musculus" or "homo_sapiens". This read count data has been processed using the iRAP pipeline (Fonseca *et al.*, 2014) which includes quality control, alignment to the reference genome using TopHat2 (Kim *et al.*, 2013), and quantification of the mapped reads per gene using HTSeq (Anders *et al.*, 2015). Because we are focusing here on bulk RNA-seq data we excluded single-cell data from the dataset. This resulted in 9,421 samples for human and 12,787 for mouse. After inspection of the distribution of total number of reads per sample, human samples with less than 2.5 million reads and mouse samples with less than 3 million reads were filtered out. Finally, cell type and tissues with less than 20 samples were removed from the further analysis. The final two datasets contained 8,796 human and 12,114 mouse samples, produced by 401 and 630 studies, covering 68 human and 76 mouse cell types and tissues, respectively (Supplementary Tables S1 and S2).

#### Evaluation of Correlation Network Quality

The processing steps described in the main paper resulted in 7,200 (3,400 human and 3,800 mouse) sets of genome-wide cell type or tissue-specific gene co-expression predictions, which we refer to as "co-expression networks". The main goal of this study is to gain understanding into what are the critical features that distinguish good co-expression networks from bad ones. Because there is no gold standard co-expression network available, we first defined eight measures of quality based on the enrichment of biologically meaningful features (functional annotations of genes and regulatory DNA motifs in promoter sequences) among co-expressed genes. To facilitate comparison between networks, these eight measures were finally combined into a single quality score (see description below).

For each gene $X$ in a co-expression network, we define $set_X$ to be the 100 genes with the highest correlation of expression with $X$ (excluding $X$ itself). Our quality measures are based on the enrichment of biologically meaningful features among the genes in $set_X$. These measures should not be interpreted as strict measures of accuracy of for example functional annotation predictions, but rather as rough indicators of quality of the inferred gene co-expression values.

**GO enrichment frequency:** Genes involved in the same biological process are expected to be co-expressed more frequently than unrelated sets of genes. In a high-quality co-expression network, we would expect the genes in $set_X$ to share a functional annotation (Figure 1B). In contrast, in a low-quality network (e.g. a randomly generated network) we expect genes in $set_X$to have a random set of annotations. We define $Enrichment_{MF}$, $Enrichment_{BP}$, and $Enrichment_{CC}$ as the fraction of genes in a network for which $set_X$ contained one or more significantly enriched GO terms (after correction for multiple testing) for Molecular Function (MF), Biological Process (BP) and Cellular Component (CC) GO terms.

**GO enrichment accuracy:** Where we found $set_X$ to have enriched GO terms, we checked if the enriched terms overlapped with the GO terms of gene $X$. $Accuracy_{MF}$, $Accuracy_{BP}$, and $Accuracy_{CC}$ were defined as the fraction of genes in the network for which this was the case for MF, BP, and CC GO terms.

**TFBS enrichment frequency:** Genes with similar expression profiles are likely to be under the control of a shared regulatory mechanism, including regulation by a similar set of transcription factors (TFs). In a high-quality co-expression network, we would therefore expect the genes in $set_X$ to contain a shared set of transcription factor binding sides (TFBSs). We define $Enrichment_{TFBS}$ as the fraction of genes in a network for which the promoter sequences of $set_X$ contained one or more significantly enriched TFBSs.

**TFBS enrichment accuracy:** Where we found $set_X$ to have enriched TFBSs, we checked if the promoter of gene $X$ contained one or more of those TFBSs. $Accuracy_{TFBS}$ was defined as the fraction of genes in the network for which this was the case.

Correlation between the eight quality measures was high (range 0.60 to 0.98). Principal component analysis (PCA; after standardizing each quality measure to mean 0 and standard deviation 1) revealed that 81.4% of the total variation in the eight quality measures could be explained by the first principal component (PC; Supplementary Figure S1A). We decided to use this first PC as the general quality score, $Quality$, after rescaling to the range 0 to 1. The correlation between $Quality$ and each of the quality measures was high (range 0.77 to 0.96; Supplementary Figure S1B-C). In conclusion, $Quality$ captures well the general trend of the eight quality measures and simplifies the comparison between the quality of different networks.

#### Gene Ontology analysis

Gene-to-GO term association data was obtained from the Mouse Genome Informatics website (http://www.informatics.jax.org/) for mouse, and from the EBI database (ftp://ftp.ebi.ac.uk/pub/databases/GO/goa/HUMAN/goa_human.gaf.gz) for human. The basic version of the Gene Ontology (go-basic.obo) was obtained from GO Consortium website (http://geneontology.org/).

GO terms were mapped to their parent terms upward in the GO graph structure. Genes (Entrez IDs) assigned to a particular GO term were also assigned to that term's parent terms. GO term enrichment in sets of 100 correlated genes (see above) was evaluated using hypergeometric tests: for every GO term the total number of genes associated with the term was compared with the number of genes in the set of 100 genes (Ensembl ids converted to Entrez ids), and a p-value was estimated using a hypergeometric distribution. The Bonferroni correction was used to adjust p-values for multiple testing, and corrected p-values < 0.01 were regarded as significant.

#### Transcription Factor Binding Site Analysis

Position Weight Matrices (PWMs) were obtained from the JASPAR database (JASPAR_CORE redundant vertebrate PWMs, version of October 2016; 635 PWMs in total) (Mathelier *et al.*, 2016). Promoter sequences (region -500 to +200 around transcription start sites) for all human (hg19/GRCh37) and mouse (mm10/GRCm38) Refseq genes were downloaded using the UCSC Table Browser (Karolchik *et al.*, 2004). We also extracted 10,000 randomly selected regions of the human genome of length 2kb and used them to set a threshold score for each PWM. Threshold scores were set so that each PWM would return on average 1 hit per 5,000 base pairs. A threshold score could be set for 618 PMWs (17 PWMs failed because of low information content).

Vertebrate promoters can be roughly divided into two classes: CpG island-associated promoters and non-CpG island promoters (Lenhard *et al.*, 2012; Illingworth and Bird, 2009). CpG island-associated promoters have on average a higher GC content. To avoid biases caused by differences in GC content and CpG scores, we conducted PWM enrichment analysis as described before (Vandenbon *et al.*, 2016). In brief, we classified all human and all mouse promoter sequences into two classes: promoters with high GC content and high CpG scores, and promoters with low GC content and low CpG scores. For each PWM $p$, we calculated $fr_{p,high}$, the fraction of high GC content promoters that contain a hit for $p$. Similarly, we calculated $fr_{p,low}$, the fraction of low GC content promoters that contain a hit for $p$. For the prediction of enriched PWM motifs in a set of promoters $D$, we counted $h_{p,D}$, the number of sequences that contain a hit for $p$, as well as the number of sequences in $D$ that were classified in the high GC content class ($n_{high}$) and in the low GC content class ($n_{low}$), respectively. Finally, using a binomial distribution, we calculated the probability of observing $h_{p,D}$ or more hits for $p$ in a set of $n_{high}$ high GC content and $n_{low}$ low GC content sequences, given $fr_{p,high}$ and $fr_{p,low}$. This probability was corrected for multiple testing using the Bonferroni correction, and PWMs with a corrected p value < 0.01 were considered as significantly enriched in the input set $D$.

#### Supplementary Tables

| | **tissue or cell type** | **no. of samples** | **no. of genes in co-expression network** | **included in validation set?** |
|---|---|---|---|---|
| 1 | fibroblast | 645 | 21,070 | Yes |
| 2 | ileum | 623 | 18,655 | |
| 3 | lymphoblastoid | 481 | 16,687 | |
| 4 | breast cancer | 427 | 27,783 | |
| 5 | acute myeloid leukemia | 402 | 24,454 | |
| 6 | PBMC | 376 | 20,579 | |
| 7 | lung | 321 | 22,531 | Yes |
| 8 | liver | 292 | 22,937 | |
| 9 | epithelial cell | 268 | 21,796 | |
| 10 | brain | 244 | 27,688 | |
| 11 | pancreas | 228 | 21,400 | Yes |
| 12 | colon | 210 | 20,354 | Yes |
| 13 | B cell | 199 | 21,648 | Yes |
| 14 | monocyte | 193 | 19,557 | |

| | | | | |
|---|---|---|---|---|
| 15 | induced pluripotent stem cell | 187 | 20,283 | |
| 16 | skeletal muscle | 170 | 18,775 | |
| 17 | embryonic stem cell | 169 | 22,967 | |
| 18 | macrophage | 169 | 22,758 | Yes |
| 19 | prostate | 163 | 25,482 | |
| 20 | colon cancer | 150 | 21,036 | |
| 21 | bone marrow | 143 | 24,409 | |
| 22 | CD4 T cell | 135 | 22,510 | |
| 23 | lung cancer | 135 | 20,883 | |
| 24 | prefrontal cortex | 129 | 26,077 | |
| 25 | heart | 113 | 19,468 | |
| 26 | breast | 111 | 21,977 | |
| 27 | neural progenitor cell | 98 | 24,969 | Yes |
| 28 | melanoma | 97 | 22,832 | |
| 29 | placenta | 86 | 19,210 | |
| 30 | embryonic kidney | 74 | 21,715 | |
| 31 | neuron | 72 | 25,675 | |
| 32 | granulocyte | 70 | 19,814 | |
| 33 | osteosarcoma | 67 | 16,838 | |
| 34 | adipose tissue | 66 | 19,435 | |
| 35 | kidney | 62 | 23,379 | Yes |
| 36 | MCF-7 | 59 | 20,334 | |
| 37 | endothelial cell | 58 | 21,238 | |
| 38 | HeLa | 56 | 22,011 | |
| 39 | CD8 T cell | 51 | 19,860 | |
| 40 | ovarian cancer | 51 | 24,595 | |
| 41 | cervical cancer | 50 | 23,836 | Yes |
| 42 | lymphoma | 49 | 28,063 | |
| 43 | hepatocyte | 48 | 15,570 | Yes |
| 44 | neutrophil | 46 | 19,667 | |
| 45 | salivary gland | 46 | 29,817 | |
| 46 | HUVEC | 45 | 21,110 | |
| 47 | keratinocyte | 45 | 20,350 | |
| 48 | frontal cortex | 44 | 18,232 | |
| 49 | mammary gland | 44 | 25,746 | |
| 50 | thyroid | 43 | 23,415 | Yes |
| 51 | esophagus | 41 | 20,439 | Yes |
| 52 | ovary | 38 | 23,973 | |
| 53 | osteoblast | 37 | 16,689 | Yes |
| 54 | pluripotent stem cell | 35 | 17,358 | |

| | | | | |
|---|---|---|---|---|
| 55 | prostate cancer | 29 | 23,663 | |
| 56 | cerebellum | 28 | 20,920 | |
| 57 | dendritic cell | 28 | 21,994 | |
| 58 | retina | 28 | 18,675 | |
| 59 | thyroid cancer | 28 | 25,167 | |
| 60 | CLL | 27 | 19,035 | |
| 61 | sarcoma | 27 | 24,168 | Yes |
| 62 | adipocyte | 26 | 14,258 | |
| 63 | mesenchymal stem cell | 25 | 18,070 | Yes |
| 64 | white adipose tissue | 24 | 16,488 | Yes |
| 65 | natural killer cell | 23 | 19,389 | |
| 66 | cancer | 21 | 26,083 | |
| 67 | cerebral cortex | 21 | 20,773 | |
| 68 | neocortex | 20 | 16,623 | Yes |

**Supplementary Table S1: Human datasets.** The cell type or tissue, the number of RNA-seq samples, and the number of genes included in the final co-expression network is shown. The last column indicates which datasets were included in the validation set.

| | **tissue or cell type** | **no. of samples** | **no. of genes in co-expression network** | **included in validation set?** |
|---|---|---|---|---|
| 1 | liver | 2,644 | 16,287 | |
| 2 | embryonic stem cell | 742 | 20,955 | |
| 3 | embryonic fibroblast | 562 | 17,864 | |
| 4 | macrophage | 556 | 17,977 | Yes |
| 5 | brain | 518 | 20,893 | Yes |
| 6 | heart | 507 | 18,347 | |
| 7 | hippocampus | 460 | 19,488 | Yes |
| 8 | cerebellum | 431 | 20,472 | |
| 9 | cortex | 317 | 19,656 | |
| 10 | spleen | 266 | 19,057 | |
| 11 | lung | 241 | 19,246 | |
| 12 | mammary gland | 221 | 19,174 | |
| 13 | testis | 216 | 23,122 | |
| 14 | B cell | 202 | 20,650 | |
| 15 | CD4 T cell | 199 | 17,300 | Yes |
| 16 | neuron | 161 | 21,232 | |
| 17 | kidney | 152 | 19,643 | |

| | | | | |
|---|---|---|---|---|
| 18 | whole embryo | 149 | 15,399 | Yes |
| 19 | retina | 140 | 18,522 | |
| 20 | cardiomyocyte | 138 | 17,309 | |
| 21 | Th17 | 122 | 15,888 | |
| 22 | white adipose tissue | 116 | 19,900 | |
| 23 | CD8 T cell | 110 | 16,718 | Yes |
| 24 | thymus | 102 | 21,358 | Yes |
| 25 | muscle | 100 | 16,188 | |
| 26 | spinal cord | 99 | 19,828 | |
| 27 | hematopoietic stem cell | 98 | 21,466 | |
| 28 | microglia | 98 | 19,901 | |
| 29 | fibroblast | 95 | 19,727 | Yes |
| 30 | dendritic cell | 90 | 17,107 | |
| 31 | neocortext | 75 | 14,482 | |
| 32 | neural stem cell | 75 | 17,587 | |
| 33 | iPS | 73 | 18,721 | Yes |
| 34 | prefrontal cortex | 71 | 16,031 | Yes |
| 35 | Treg | 71 | 19,567 | |
| 36 | colon | 70 | 18,570 | |
| 37 | cerebral cortex | 62 | 17,569 | |
| 38 | dorsal root ganglion | 61 | 17,996 | |
| 39 | pro-B | 60 | 15,496 | Yes |
| 40 | myoblast | 59 | 17,415 | |
| 41 | pre-B | 56 | 16,083 | |
| 42 | spermatocyte | 55 | 19,841 | |
| 43 | brown adipose tissue | 54 | 16,610 | |
| 44 | osteoblast | 53 | 15,999 | Yes |
| 45 | frontal cortex | 49 | 18,596 | |
| 46 | motoneuron | 48 | 16,020 | |
| 47 | trophoblast stem cell | 47 | 15,389 | |
| 48 | AML | 44 | 16,143 | |
| 49 | astrocyte | 44 | 20,342 | |
| 50 | spermatid | 43 | 19,840 | |
| 51 | endothelial cell | 42 | 20,169 | |
| 52 | midbrain | 38 | 19,159 | |
| 53 | thymocyte | 38 | 14,956 | |
| 54 | neural progenitor cell | 37 | 18,891 | Yes |
| 55 | Th0 | 37 | 14,564 | |
| 56 | small intestine | 36 | 17,929 | Yes |
| 57 | pancreas | 35 | 16,682 | |

| | | | | |
|---|---|---|---|---|
| 58 | skeletal muscle | 35 | 19,382 | |
| 59 | hepatocyte | 32 | 14,696 | Yes |
| 60 | 3T3 | 31 | 15,091 | Yes |
| 61 | forebrain | 29 | 20,229 | |
| 62 | placenta | 29 | 16,515 | |
| 63 | large intestine | 28 | 19,992 | |
| 64 | duodenum | 27 | 30,937 | |
| 65 | ileum | 27 | 18,358 | |
| 66 | oocyte | 27 | 14,937 | Yes |
| 67 | uterus | 26 | 15,813 | |
| 68 | intestine | 25 | 19,746 | Yes |
| 69 | Th1 | 25 | 17,516 | |
| 70 | thyroid | 24 | 12,941 | Yes |
| 71 | satellite cell | 23 | 17,247 | |
| 72 | stomach | 23 | 19,209 | |
| 73 | mesoderm | 22 | 16,219 | |
| 74 | olfactory bulb | 21 | 17,698 | |
| 75 | trophoblast | 21 | 13,038 | |
| 76 | medullary thymic epithelial cell | 20 | 22,097 | |

**Supplementary Table S2: Mouse datasets.** The cell type or tissue, the number of RNA-seq samples, and the number of genes included in the final co-expression network is shown. The last column indicates which datasets were included in the validation set.

#### Supplementary Figures

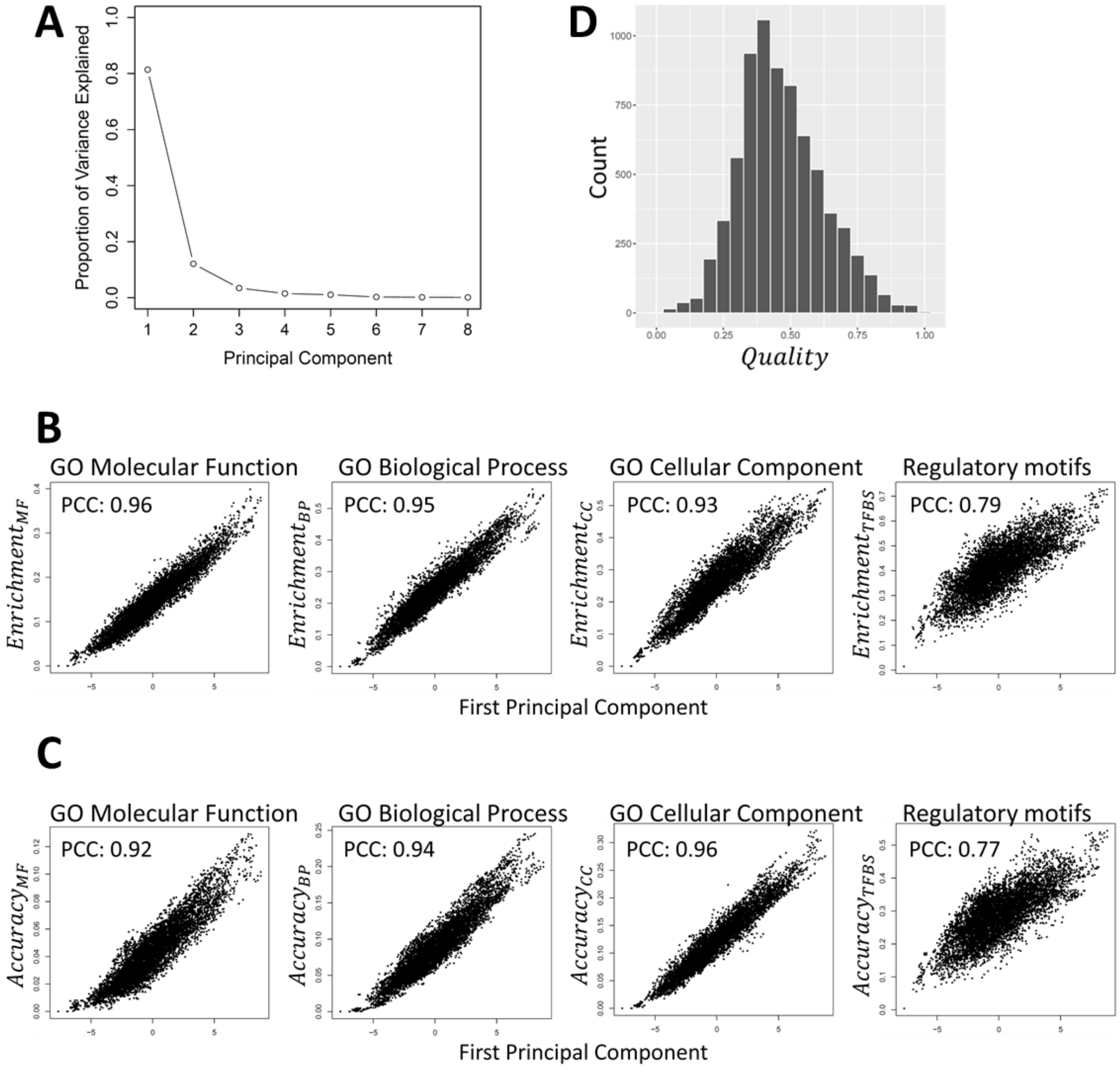

**Supplementary Figure S1: Principal Component Analysis of the eight quality measures. (A)** Proportion of the variance in the eight quality measures explained by the principal components (PCs). The first PC explains 81.4% of the total variance. **(B-C)** Scatterplots of PC1 (X-axis) versus each of the eight individual quality measures (Y-axes). Each plot shows 7,200 dots, each representing a genome-wide gene-gene co-expression network for a cell type or tissue. The Pearson correlation coefficient (PCC) is indicated in each plot. **(B)** The first PC versus measures for enrichment frequency, and **(C)** versus measures for enrichment accuracy. **(D)** Distribution of the 7,200 general quality scores, $Quality$.

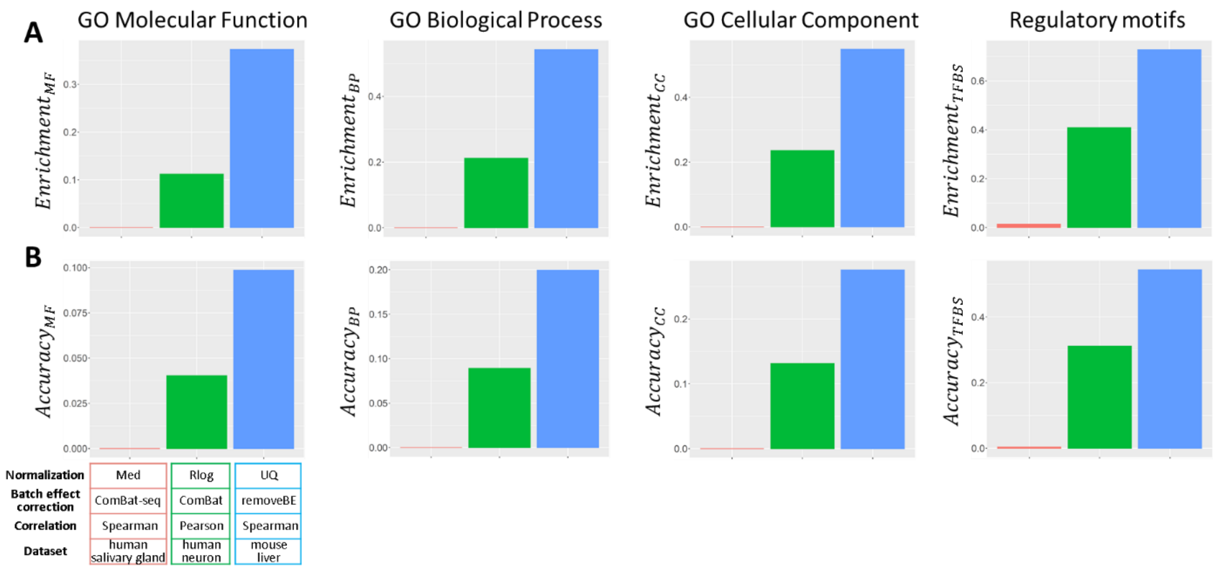

**Supplementary Figure S2: Raw quality measures of three networks.** This figure shows the remaining 8 raw quality measures of the worst (red), the median (green), and the best (blue) network for GO Molecular Function, Biological Process, Cellular Component and Regulatory motifs in promoter sequences. **(A)** shows the frequency of enrichment and **(B)** the accuracy.

**(next page) Supplementary Figure S3: Overview of the performance of each workflow on each of the 144 datasets.** For each of the 50 workflows the relative performance on each of the 144 datasets is visualized. Workflows are ordered in order of average overall performance as in Figure 2D in the main text. Datasets are ordered according to sample counts. Colors indicate the relative performances of the 50 workflows on each dataset.

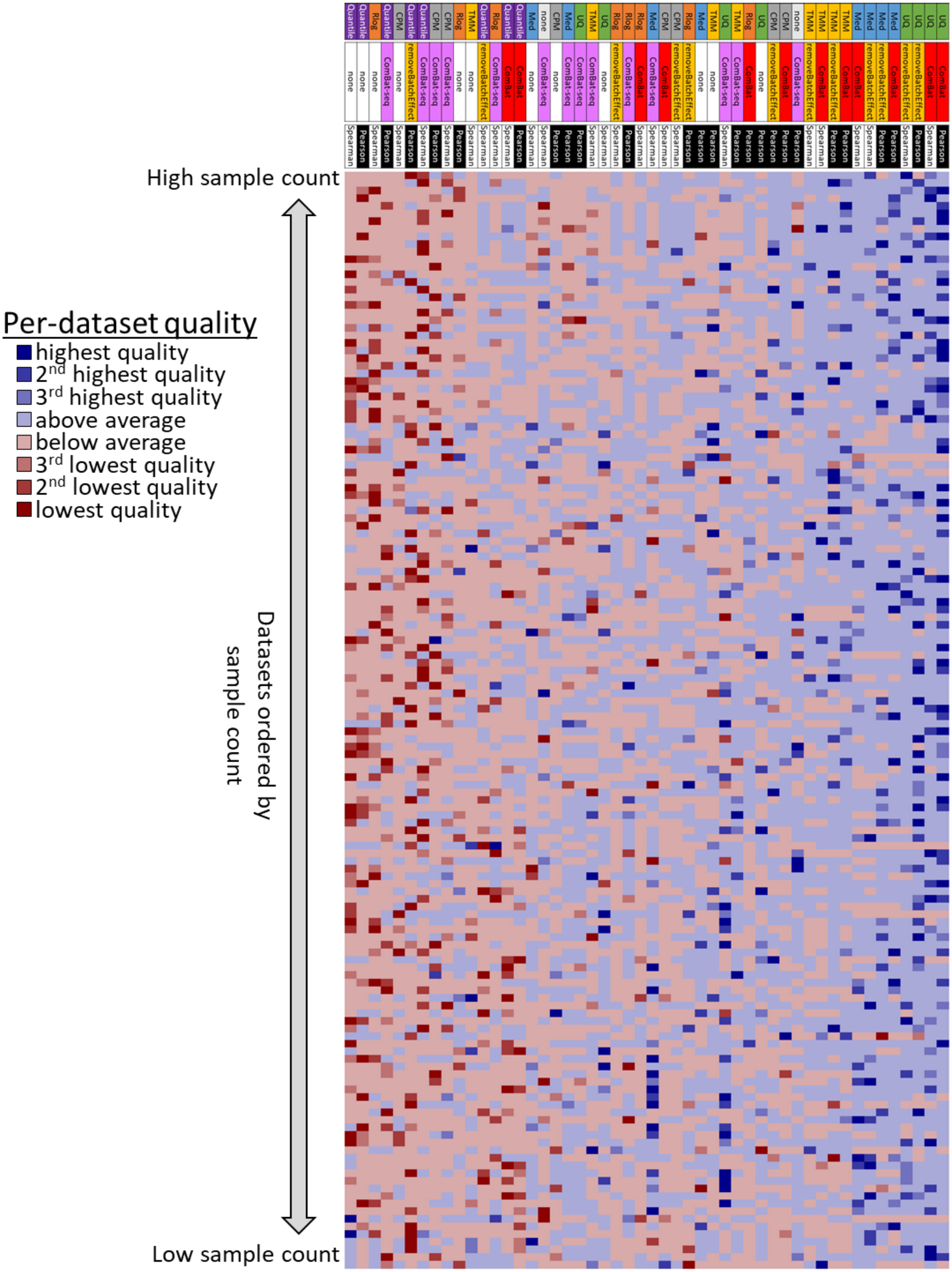
High sample count
Low sample count
Datasets ordered by sample count
Per-dataset quality
highest quality
2nd highest quality
3rd highest quality
above average
below average
3rd lowest quality
2nd lowest quality
lowest quality
